# CellMethMarker provides an integrated database of human cell- and tissue-type-specific DNA methylation markers

**DOI:** 10.64898/2026.10.23.760902

**Authors:** Xiaojun Sun, Yujie Peng, Yingyu Yi, Lina Fu, Fang Liu, Yali Xie, Qing Dai, Hongcang Gu, Jiantao Shi, Hongzhi Wang, Fan Zhang

## Abstract

Cell- and tissue-type-specific DNA methylation markers provide epigenetic signatures of cellular identity and are widely used to infer cellular origin from bulk and single-cell methylomes. Existing resources mainly emphasize disease-associated biomarkers, methylome atlases, genomic annotations or RNA-based cell markers, leaving an unmet need for an integrated region-level resource dedicated to such markers. Here we present CellMethMarker (http://gu-zhanglab.hfcas.ac.cn/CellMethMarker), a curated database integrating 347,514 *de novo* markers derived from uniformly processed whole-genome and reduced-representation bisulfite sequencing datasets, 2,203,507 literature-derived marker records and harmonized metadata for 2,810 samples. CellMethMarker enables cell- and tissue-centered marker retrieval, coordinate-based search, genome-browser visualization, interactive methylation analysis and data download. In an independent single-cell methylome atlas, these markers recover the expected cell-type-associated methylation patterns, and a platform-matched reference built from them lowers the mean deconvolution error from 0.365 to 0.144. CellMethMarker therefore links region-level DNA methylation variation to cellular identity and supports methylome-based cell-type annotation and deconvolution.

## Introduction

DNA methylation is a key epigenetic modification involved in transcriptional regulation, cell identity maintenance, and disease processes such as cancer and developmental disorders (Greenberg and Bourc’his 2019; El Omari et al. 2025). Extensive studies have demonstrated that human cell types possess distinct DNA methylation signatures, often referred to as cell-type-specific methylation markers (Slieker et al. 2013; Moss et al. 2018). These markers, commonly represented as differentially methylated positions (DMPs), differentially methylated regions (DMRs) or differentially methylated blocks (DMBs), provide powerful tools for investigating lineage specification, cellular deconvolution, and epigenetic regulation (Lehmann-Werman et al. 2016; Moss et al. 2020; Fox-Fisher et al. 2021, 2023). Profiling technologies, including whole-genome bisulfite sequencing (WGBS), reduced representation bisulfite sequencing (RRBS) and array-based assays such as the Illumina 450K and EPIC arrays, have greatly facilitated the discovery of cell- and tissue-type-specific markers across diverse biological contexts (Slieker et al. 2013; Caggiano et al. 2021; Li et al. 2023; Loyfer et al. 2023).

Existing resources capture important aspects of methylation biology but do not integrate the region-level markers needed to discriminate human cell and tissue types across studies and platforms. MethHC 2.0 (Huang et al. 2021), DNMIVD (Ding et al. 2020), DiseaseMeth 3.0 (Xing et al. 2022), and MethMarkerDB (Zhu et al. 2024) primarily catalog disease- or cancer-associated methylation changes; MethyMark (Liu et al. 2016), NGSmethDB (Lebrón et al. 2017), scMethBank (Zong et al. 2022), and MethBank 4.0 (Zhang et al. 2023) provide methylome atlases or genomic annotations; and CellMarker 2.0 (Hu et al. 2023) and Cell Taxonomy (Jiang et al. 2023) catalog RNA-derived or multimodal cell markers. These resources are valuable for their intended purposes, but none was designed to unify curated literature evidence, uniformly called DNA methylation markers and sample-level context for human cell- and tissue-type annotation.

In practice, literature-derived markers remain dispersed across publications, are reported in different genome assemblies, and are defined at heterogeneous molecular resolutions, whereas public WGBS and RRBS datasets require standardized processing before cross-study comparison. Cell and tissue labels are free text, so the same population appears under several names. An integrated resource is therefore needed to combine curated literature evidence, uniformly called markers, harmonized sample metadata, ontology-based cell and tissue annotations, and tools for genomic and cell-origin interpretation.

To address this gap, we developed CellMethMarker, a curated and web-browsable resource for human cell- and tissue-specific DNA methylation markers. CellMethMarker integrates four connected data layers: a *de novo* atlas of 347,514 cell- and tissue-specific marker records generated from uniformly processed WGBS and RRBS datasets, a manually curated collection of 2,203,507 literature-derived methylation marker records from 78 eligible publications, harmonized metadata for 2,810 human methylation samples, and 2,171 processed sample-level methylation profiles. These resources are standardized by genomic coordinate, assay type, cell and tissue annotation, and ontology identifier, and are connected through a web interface that supports custom marker search, literature marker search, sample metadata search, coordinate search, genome browser visualization, methylation heatmap and sample clustering analysis, data download, and help documentation. We further demonstrate the utility of CellMethMarker through two application-oriented case studies: methylation-based cell-type annotation in an independent human single-cell methylome atlas, and construction of a platform-matched RRBS reference atlas that improves methylation deconvolution with the fragment-level UXM algorithm (Loyfer et al. 2023). Together, CellMethMarker provides an integrated framework for linking DNA methylation markers to cell and tissue identity, sample context, genomic location, and downstream cell-origin interpretation.

## Results

### A resource linking methylation markers to their experimental context

CellMethMarker is organized as an integrated resource for searching, visualizing, and applying human cell- and tissue-specific DNA methylation markers (Fig. 1). At its core are four interconnected data layers that span the full path from marker discovery to sample context. Specifically, the database comprises (i) an atlas of 347,514 *de novo* custom markers generated from uniformly processed WGBS and RRBS datasets; (ii) 2,203,507 manually curated literature-derived methylation marker records from 78 eligible publications; (iii) harmonized metadata describing the human methylation samples underlying or related to these marker resources, and (iv) 2,171 processed sample-level methylation profiles (Fig. 1A, B).

**Fig. 1:**
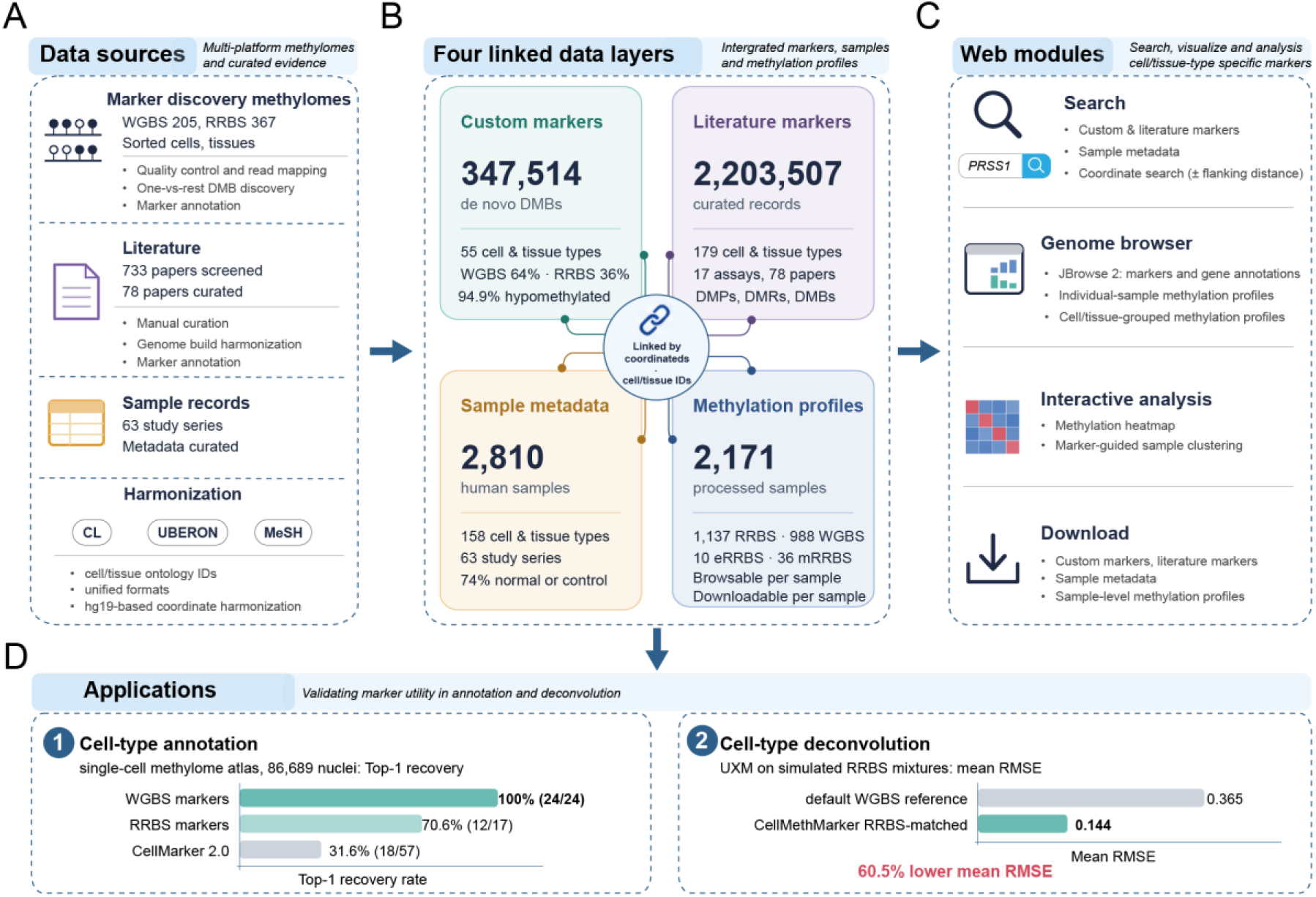
Organization and applications of CellMethMarker. **A** Data sources and processing workflow. Whole-genome bisulfite sequencing (WGBS) and reduced-representation bisulfite sequencing (RRBS) methylomes are processed to identify custom markers; literature markers are manually curated from published studies; and sample metadata are collected from study records. Data are harmonized using cell/tissue ontology identifiers and standardized formats. hg19 was used as the primary genome assembly for online search and visualization, with an hg38 liftOver version of literature-derived markers also provided for download. **B** Four linked data layers comprise 347,514 custom markers, 2,203,507 literature-derived marker records, metadata for 2,810 human samples, and processed methylation profiles from 2,171 samples. These layers are connected through genomic coordinates and ontology identifiers, with summary statistics shown for each layer. **C** Web modules support marker and sample metadata searches, coordinate searches with a user-defined flanking distance, and genome-browser visualization of marker regions alongside individual-sample and cell/tissue-grouped methylation profiles using JBrowse 2. Additional functions include interactive methylation heatmaps, marker-guided sample clustering, and data download. **D** Applications demonstrate cell-type annotation using a single-cell methylome atlas containing 86,689 nuclei and cell-type deconvolution using simulated RRBS mixtures. Annotation performance is summarized by Top-1 recovery, whereas deconvolution performance is measured by root mean square error (RMSE) averaged across benchmark cell and tissue types.

These layers are linked through standardized genomic coordinates, assay information, cell and tissue labels, ontology identifiers, gene annotations and literature evidence, enabling users to move from a marker region to its biological context and supporting evidence. To make these resources accessible, CellMethMarker provides dedicated modules for custom marker search, literature marker search, sample metadata search, coordinate search, genome browser visualization, methylation heatmap and sample clustering analysis, and data download (Fig. 1C). Cell-type identity recovery was evaluated in an independent human single-cell methylome atlas comprising 86,689 nuclei (Zhou et al. 2026). Cell-type deconvolution was assessed using UXM (Loyfer et al. 2023) on simulated RRBS mixtures, comparing its default WGBS-derived reference with an RRBS-matched reference built from CellMethMarker custom markers (Fig. 1D). Together, these components form a web-browsable framework that connects methylation marker regions with cell and tissue identity, genomic location, sample context, and downstream methylome interpretation.

### Resource coverage and assay-specific marker collections

The custom-marker atlas was built from 367 RRBS samples representing 27 cell types and 205 WGBS samples representing 34 cell types that had passed quality control and were included in one-versus-rest marker discovery. Importantly, these discovery samples were enriched for sorted cell populations and dissected tissue specimens, favoring cell- and tissue-intrinsic methylation signals over signals dominated by heterogeneous bulk composition. The analysis identified 347,514 custom DMB records covering 55 cell and tissue types, comprising 223,505 WGBS-derived (64.3%) and 124,009 RRBS-derived (35.7%) markers (Fig. 2A). Of these, 329,784 (94.9%) were hypomethylated in the cell type of interest relative to all others (Supplementary Fig. S1A).

**Fig. 2:**
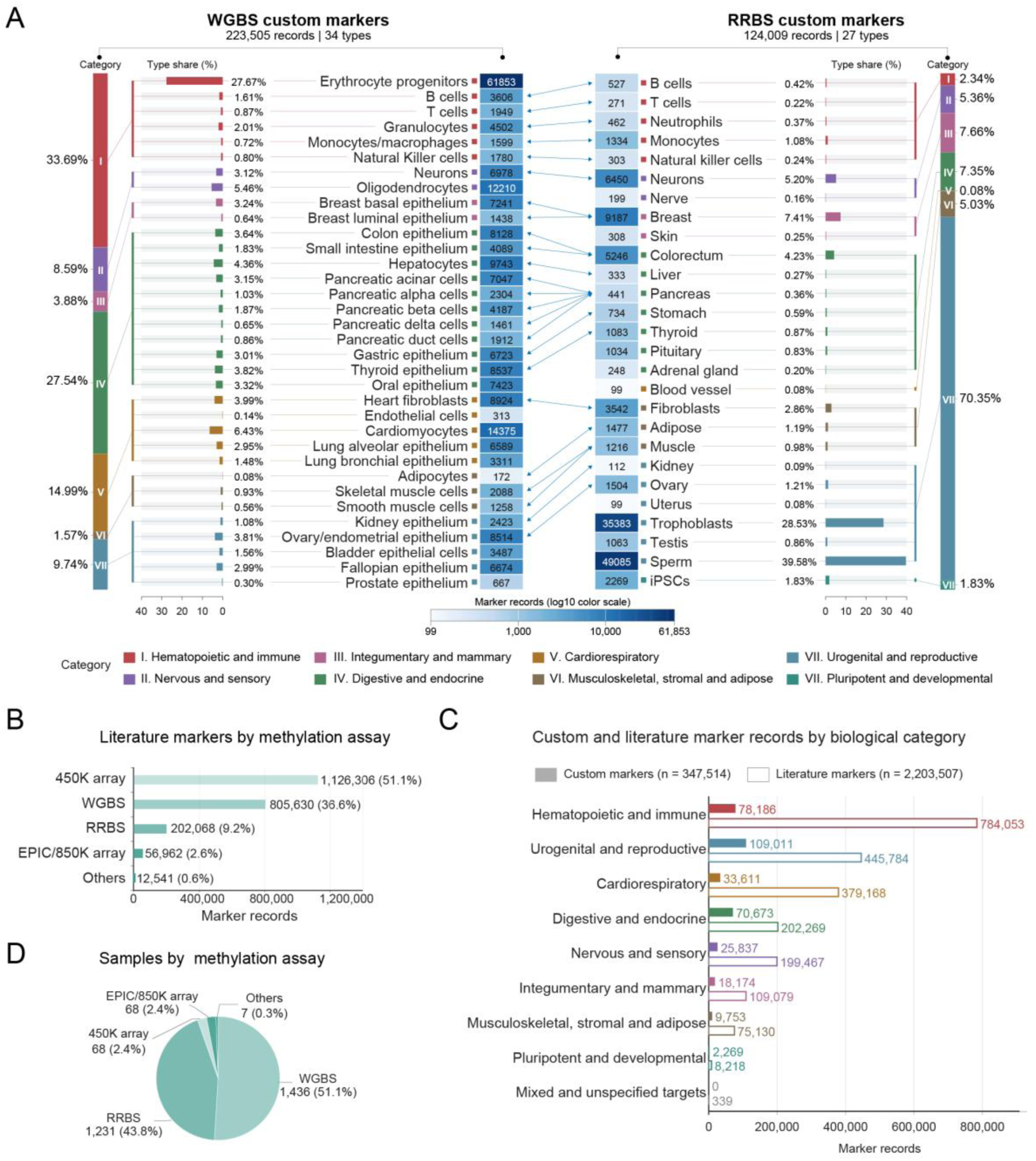
Coverage of the resource. **A** The left and right panels summarize custom marker records derived from whole-genome bisulfite sequencing (WGBS; 223,505 records across 34 cell/tissue types) and reduced-representation bisulfite sequencing (RRBS; 124,009 records across 27 types), respectively. For each sequencing method, the vertical stacked bar labeled Category shows the proportion of marker records assigned to each broad biological category, with colors and Roman numerals corresponding to the categories listed in the legend. Adjacent horizontal bars labeled Type share (%) indicate the contribution of each individual cell or tissue type to the total marker count for that sequencing method. The central heatmap-style count columns display marker-record counts using a shared log10 color scale, with exact counts shown in each cell. Colored guide lines link each cell or tissue type to its corresponding broad biological category, and double-headed arrows connect biologically related annotations between the WGBS and RRBS marker collections. Counts represent marker records without additional deduplication by genomic coordinates. **B** Literature-derived marker records by methylation assay. Assay types outside the categories shown are pooled as “Others”. **C** Distribution of custom and literature-derived methylation marker records across biological categories. Solid bars represent custom markers pooled from WGBS and RRBS datasets (347,514 records), and outlined bars represent literature-derived markers (2,203,507 records). Colors denote nine broad biological categories, ordered by the combined record count from both collections. Bar lengths indicate marker-record counts on a linear scale, with exact counts shown at the bar ends. Counts are reported at the record level, without additional deduplication by genomic coordinates. **D** Curated samples by methylation assay. WGBS, whole-genome bisulfite sequencing; RRBS, reduced-representation bisulfite sequencing; 450K array, Illumina Infinium HumanMethylation450 BeadChip; EPIC/850K array, Illumina Infinium MethylationEPIC BeadChip.

Marker representation varied substantially across cell and tissue types and between sequencing platforms (Fig. 2A). Erythrocyte progenitors contributed the largest WGBS-derived marker set (61,853 records, 27.7%), whereas sperm (49,085 records, 39.6%) and trophoblasts (35,383 records, 28.5%) together accounted for 68.1% of RRBS-derived markers. Grouping cell and tissue types into eight broad biological categories further highlighted differences in their representation across the two platforms (Methods), while revealing shared biological targets that could be directly compared between marker collections. The mean and median methylation differences (Δβ) were 0.58 and 0.59 for WGBS-derived markers and 0.64 and 0.64 for RRBS-derived markers (Supplementary Fig. S1B), indicating that the retained markers separate cell types by a wide margin rather than by marginal shifts.

CellMethMarker also contains 2,203,507 literature-derived marker records representing 179 human cell and tissue types. Of 733 screened publications, 78 met the inclusion criteria and provided extractable marker data. The curated records encompass 17 methylation assay types, dominated by 450K arrays (1,126,306 records, 51.1%), WGBS (805,630, 36.6%) and RRBS (202,068, 9.2%), and seven marker types, primarily DMPs (1,172,196, 53.2%), DMRs (963,881, 43.7%) and DMBs (52,809, 2.4%) (Fig. 2B, Supplementary Fig. S1C). Most records were originally reported in hg19 (92.7%); coordinates reported in hg18 or hg38 were converted to hg19 for online search and visualization.

Comparison across nine biological categories revealed distinct but complementary patterns of marker representation (Fig. 2C). Custom markers showed greater relative representation of urogenital and reproductive (31.4%) and hematopoietic and immune (22.5%) categories, whereas literature-derived markers were more concentrated in hematopoietic and immune (35.6%) and urogenital and reproductive (20.2%) categories. Their overlapping biological coverage provides a common framework for integrating marker evidence across data sources.

CellMethMarker contains harmonized metadata for 2,810 human DNA methylation samples from 63 study series, including the 572 samples used for custom-marker discovery. The records represent 158 cell and tissue types, 44 biological sources, seven assay types, 38 cell-sorting methods, and 45 original medical-condition labels. Blood-derived samples accounted for 49.7% of the collection; WGBS and RRBS accounted for 51.1% and 43.8% of samples, respectively (Fig. 2D); and 42.2% of samples were generated using fluorescence-activated cell sorting (FACS; Supplementary Fig. S1D). After harmonization, normal, control or non-cancer samples comprised 74.0% of the collection. Of these 2,810 samples, 2,171 had methylation profiles of sufficient quality to be reprocessed to a common format and served through the genome browser (1,137 RRBS, 988 WGBS, 10 enhanced reduced-representation bisulfite sequencing (eRRBS) and 36 multiplexed reduced-representation bisulfite sequencing (mRRBS)).

### Harmonized annotations retain source and measurement context

To support consistent interpretation across marker sources, CellMethMarker harmonizes cell and tissue annotations across custom markers, literature markers and sample metadata. Labels from different datasets and publications were manually curated, cleaned, merged when they referred to the same biological entity, and mapped to identifiers in the Cell Ontology (CL) (Diehl et al. 2016), the Uber-anatomy ontology (UBERON) (Mungall et al. 2012) or Medical Subject Headings (MeSH) using the Ontology Lookup Service and manual review (Methods). Marker records were further linked to genomic coordinates, associated genes, genomic classes, CpG-rich annotations, methylation direction, marker type, assay type and ontology identifiers, while sample metadata were organized by harmonized cell and tissue labels, biological source, assay type, study series and sample preparation information.

Because 2,042,252 of the literature-derived marker records (92.7%) were originally reported in hg19/GRCh37, CellMethMarker uses hg19 as the primary coordinate system for online search and visualization; an hg38/GRCh38 liftOver version of the literature markers is also provided for download so that the collection can be used with the current assembly. This genome-build harmonization, together with the standardized data format, facilitates consistent comparison and integration of human cell- and tissue-type-specific DNA methylation markers across experimental platforms, publications and data types.

### Comparison with existing resources

To position CellMethMarker among existing DNA methylation resources, we compared it with MethBank 4.0 (Zhang et al. 2023), scMethBank (Zong et al. 2022), and MethyMark (Liu et al. 2016) across data coverage, marker content, and analysis functionality (Table 1). MethBank 4.0 and scMethBank address scopes distinct from ours: MethBank 4.0 is a multi-species platform integrating methylomes across 23 species, and scMethBank is dedicated to single-cell DNA methylomes; neither curates cell- or tissue-type-specific markers from published studies. MethyMark shares our focus on cell-type-specific methylation markers and therefore warrants a closer comparison; CellMethMarker differs from it in kind rather than merely in scale.

**Table 1.** Comparison of CellMethMarker with existing DNA methylation resources.

| Feature | CellMethMarker | MethBank 4.0 | scMethBank | MethyMark |
| --- | --- | --- | --- | --- |
| RESOURCE SCOPE |  |  |  |  |
| **Primary focus** | Cell-type markers<sup>a</sup> | Multi-species methylomes | Single-cell methylomes | Cell-type markers<sup>a</sup> |
| **Species** | Human | 23 species | Human, mouse | Human |
| **Cell/tissue types** | 261 | 236 | 29 | 42 |
| **Data type** | Sequencing and arrays<sup>b</sup> | WGBS | Single-cell BS-seq<sup>c</sup> | WGBS |
| **Samples** | 2,810 | 1,449 | 8,328 | 50 |
| **Publication year** | — | 2023 | 2022 | 2016 |
| MARKER CONTENT & EVIDENCE |  |  |  |  |
| **Marker type** | DMB, DMR, DMP, iVMF, MHB, MHL, and CGI | DMR, curated DMGs | DMR | DMR |
| **De novo markers<sup>d</sup>** | 347,514 DMBS | 59,750,225 DMRs | 165,918 DMRs | 460,438 DMRs |
| **Literature markers** | 2,203,507 | NR | 0 | 0 |
| **De novo marker discovery** | Yes | Yes | Yes | Yes |
| **Literature marker curation<sup>e</sup>** | Yes | Yes | No | No |
| **Cross-dataset marker discovery<sup>f</sup>** | Yes | No | No | Yes |
| DATA ACCESS & ANALYSIS |  |  |  |  |
| **Sample metadata** | Yes | Yes | Yes | Yes |
| **Processed methylation download** | Yes | Yes | Yes | No |
| **Region-based marker search** | Yes | No | No | Yes |
| **Marker-guided sample clustering** | Yes | No | No | No |

Abbreviations: BS-seq, bisulfite sequencing; WGBS, whole-genome bisulfite sequencing; DMP, differentially methylated position; DMR, differentially methylated region; DMB, differentially methylated block; DMG, differentially methylated gene; iVMF, inter-individual variably methylated fragment; MHB, methylation haplotype block; MHL, methylation haplotype load; CGI, CpG island; NR, not reported or insufficiently documented. Yes, supported; No, not provided in the assessed version.
<sup>a</sup> Cell- and tissue-type-specific DNA methylation markers.
<sup>b</sup> DNA methylation sequencing and array assays, including WGBS, RRBS, 450K, EPIC, eRRBS, mRRBS, Genome-wide targeted methyl-seq.
<sup>c</sup> Bisulfite-based single-cell and single-nucleus methylation profiling, including scBS-seq, snmC-seq, scPBAT, scPBAL, scM&T-seq, scTrioSeq2, scCOOL-seq, and iscCOOL-seq.
<sup>d</sup> Counts represent DMBS for CellMethMarker, DMR records for MethBank 4.0 and scMethBank, and cell-type-associated MethyMark records for MethyMark.
<sup>e</sup> Curation of published methylation markers.
<sup>f</sup> Joint marker identification using samples from independent studies.

First, the evidence model differs: MethyMark is a single-layer *de novo* atlas of 460,438 WGBS-derived DMRs, whereas CellMethMarker maintains two complementary, provenance-tracked evidence layers — 347,514 *de novo* markers and 2,203,507 curated literature records — allowing users to distinguish markers supported by convergent evidence across studies and platforms from those reported by a single study. Second, platform and resolution coverage differ: CellMethMarker spans WGBS, RRBS, 450K and EPIC assays and marker resolutions from single positions (DMPs) to regions (DMRs) and blocks (DMBs), whereas MethyMark is restricted to WGBS-derived DMRs. Third, the context attached to each marker differs: CellMethMarker marker collections are integrated with harmonized sample metadata and available methylation profiles browsable at single-sample resolution, so that a marker region can be traced back to the samples and evidence behind it. Finally, the intended use differs: beyond retrieval, CellMethMarker connects markers to downstream analysis through coordinate-based search, genome-browser visualization and marker-guided sample clustering, as demonstrated in the cell-type annotation and deconvolution case studies. In short, MethyMark catalogs which regions are cell-type specific, whereas CellMethMarker tells users how strongly each marker is supported, in what experimental context it holds, and how it can be directly applied.

### Evidence-guided retrieval and assessment of cell-type-specific methylation markers

The four search and analysis modules described above are designed to operate as a connected workflow rather than as isolated tools. To evaluate whether the independent data layers of CellMethMarker provide mutually consistent evidence for the same biological entity, we applied this workflow to *PRSS1*, a canonical marker gene of pancreatic acinar cells in human pancreatic single-cell transcriptomic atlases (Muraro et al. 2016)(Fig. 3).

**Fig. 3:**
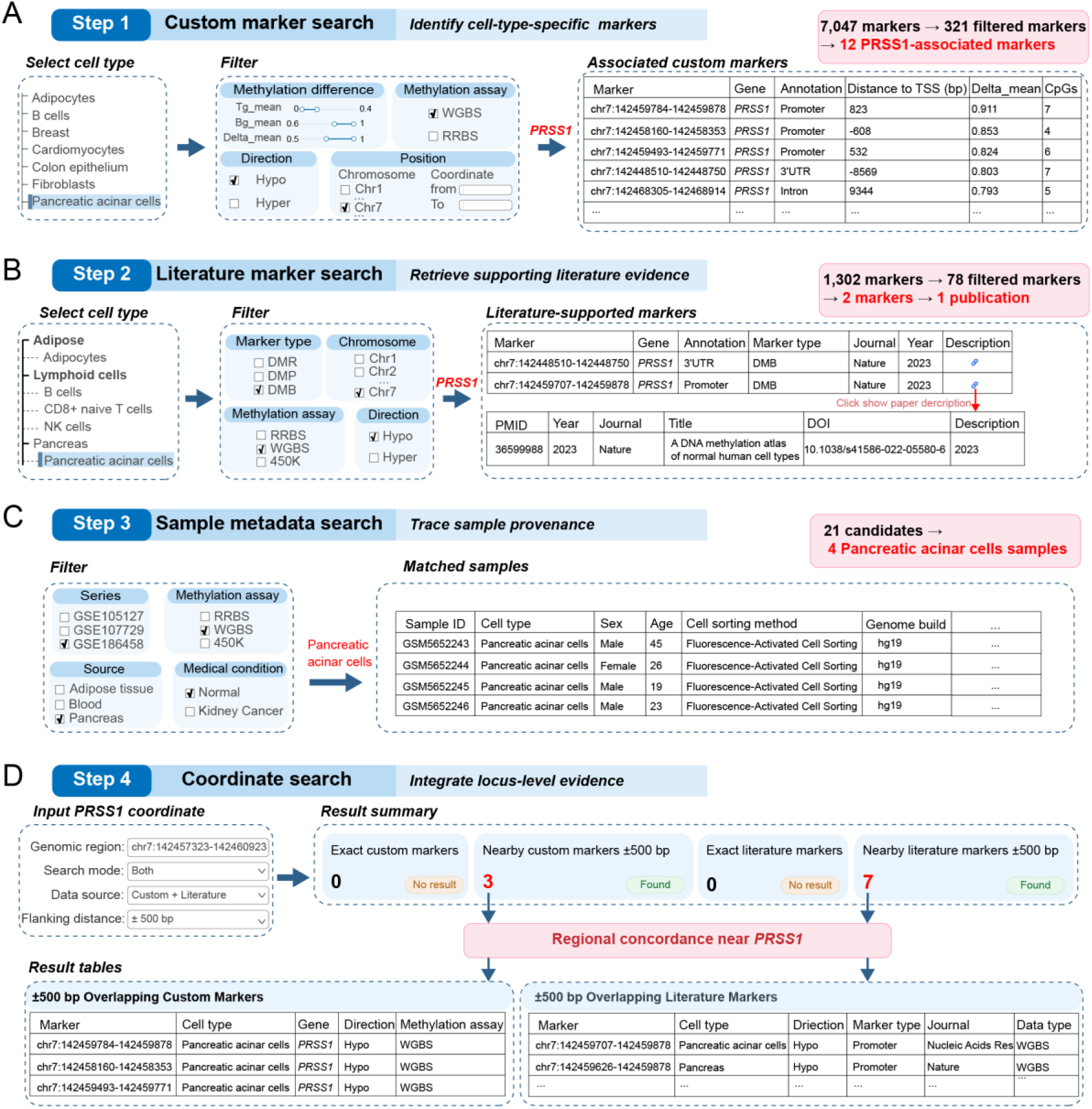
Cell-type-guided retrieval and integration of methylation marker evidence. **A** Custom marker search. Pancreatic acinar cells were selected as an example to retrieve cell-type-specific methylation markers. Markers were filtered by methylation difference, assay type, methylation direction, chromosome and genomic location, resulting in *PRSS1*-associated candidate markers with summarized genomic annotations and methylation characteristics. **B** Literature marker search. Curated literature-derived markers were queried using the same cell-type context and filtered by marker type, assay, genomic location and methylation direction. Matched *PRSS1*-associated literature markers and their corresponding publication information were displayed to provide independent supporting evidence. **C** Sample metadata search. Samples corresponding to pancreatic acinar cells were identified from curated methylation datasets by filtering study series, assay type, biological source and sample condition. The resulting sample-level metadata records provide provenance information for downstream methylation profile interpretation. **D** Coordinate search. A genomic region surrounding *PRSS1* was queried to integrate custom and literature-derived methylation markers at the locus level. Exact and nearby marker searches were summarized, and overlapping marker records from both sources were displayed to evaluate regional concordance of methylation evidence.

Custom marker search over the pancreatic acinar cell collection returned 321 candidate markers from 7,047 associated records after filtering by methylation assay, direction and difference (Δβ ≥ 0.5; Fig. 3A). Twelve of these mapped to the *PRSS1* locus; all were hypomethylated in pancreatic acinar cells relative to background cell types (Δβ = 0.501–0.911), and three overlapped promoter regions. Consistent evidence was also retrieved from the literature layer: two *PRSS1*-associated literature markers, both originating from a published WGBS atlas of normal human cell types (Loyfer et al. 2023), were retrieved together with their publication metadata (Fig. 3B), allowing direct inspection of whether a computationally derived marker has been reported before. Sample metadata search then linked these markers to four FACS-sorted pancreatic acinar cell WGBS samples with complete annotations (Fig. 3C), so that each retrieved marker can be traced to the sorted primary samples in which it was measured.

Finally, coordinate search centered on the *PRSS1* locus (chr7:142,457,323-142,460,923, ±500 bp) recovered the same three promoter-associated custom markers together with seven literature-derived markers from three independent publications (Fig. 3D); one of these studies reported *PRSS1* methylation at the pancreas tissue level, connecting tissue-level and cell-type-specific patterns. The convergence of *de novo* markers, curated literature records and sample-level provenance at a single well-characterized locus constitutes an internal cross-validation of the resource: layers built from different data types and curation strategies point to the same biological signal. Together, these connected functions turn CellMethMarker from a marker lookup table into an evidence-guided platform for discovering, evaluating and interpreting cell-type-specific methylation markers.

### Integrated marker analysis and cell-type discrimination using CellMethMarker

Beyond retrieval, CellMethMarker provides three connected analysis modules — methylation heatmap, sample clustering and genome-browser inspection — that allow users to evaluate marker behavior across samples within the platform (Fig. 4). We first examined mean WGBS methylation over the marker regions of a panel of canonical cell-identity genes across diverse human cell and tissue types (Methods). The heatmap showed the expected cell-type-restricted pattern, dominated by hypomethylation in the populations in which the corresponding genes are active (Fig. 4A) — for example, *GATA1* and *KLF1* in erythrocyte progenitors, *CD3D* and *CD3E* in T cells, *PRSS1* and *PNLIP* in pancreatic acinar cells, and *INS* and *IAPP* in pancreatic beta cells.

**Fig. 4:**
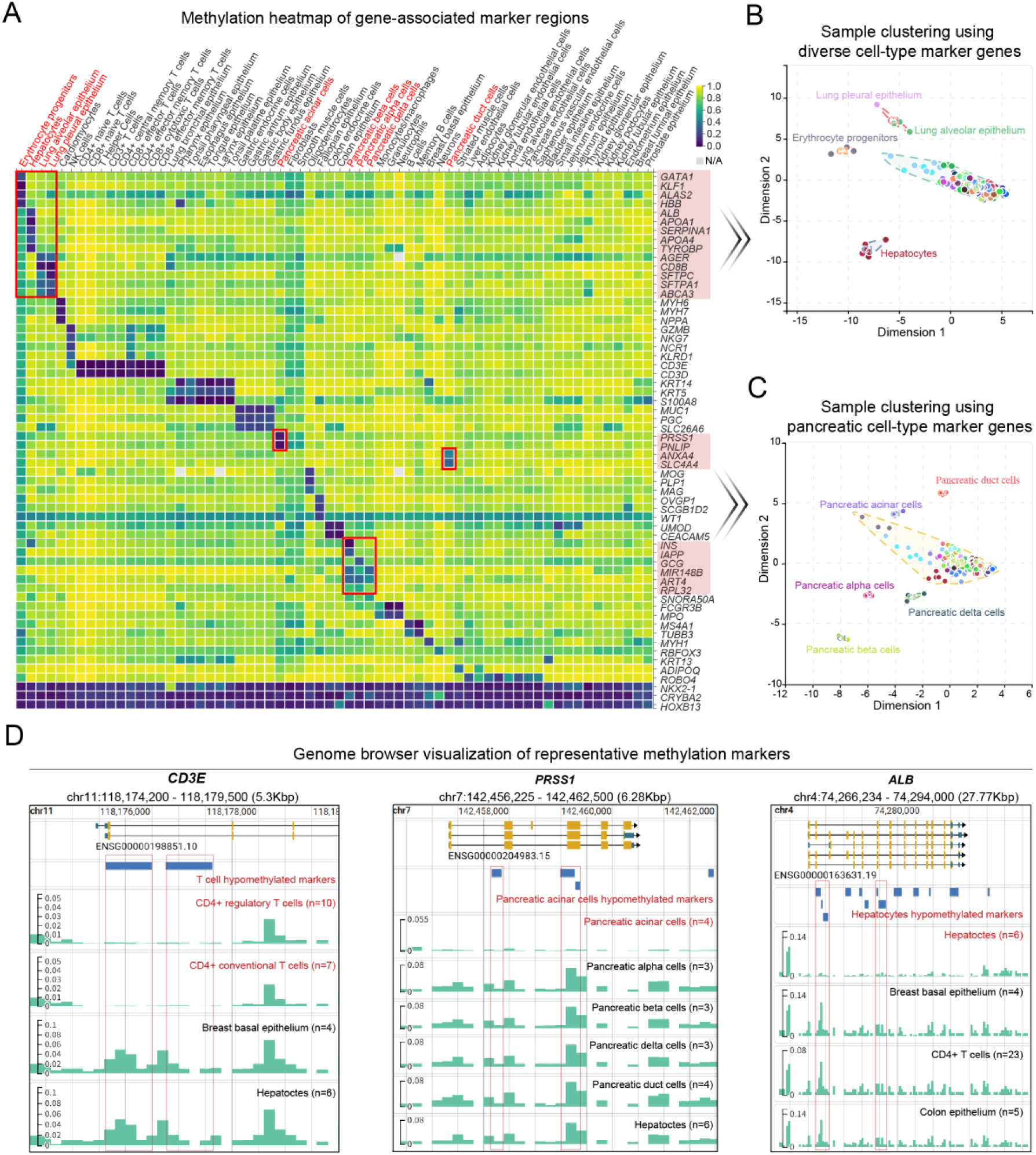
Integrated analysis of cell-type-specific DNA methylation markers using CellMethMarker. **A** Methylation heatmap of selected marker-associated genes across diverse human cell and tissue types. Rows represent genes, and columns represent cell or tissue types. Each cell shows the mean WGBS methylation level across cell-type-specific methylation marker regions associated with the corresponding gene. Colors range from purple (low methylation) to yellow (high methylation), with gray indicating unavailable data. Selected cell types and gene sets are highlighted for subsequent clustering analyses. **B** Sample clustering using marker genes exhibiting cell-type-associated hypomethylation in erythroid progenitors, hepatocytes, lung alveolar epithelium, and lung pleural epithelium. The two-dimensional projection shows the separation of samples from these cell types. **C** Sample clustering using 10 marker genes associated with five pancreatic cell types: acinar, ductal, alpha, beta, and delta cells. The resulting clusters demonstrate the ability of the selected methylation signatures to distinguish closely related cell types within the same organ. **D** Genome browser visualization of representative methylation markers associated with *CD3E*, *PRSS1*, and *ALB*. Gene annotations, cell-type-specific marker regions, and grouped WGBS methylation profiles are displayed for the corresponding genomic loci. Red boxes highlight representative hypomethylated marker regions, showing reduced methylation in the corresponding T-cell, pancreatic acinar cell, and hepatocyte populations relative to selected comparison cell types. Sample numbers for each grouped methylation track are indicated in parentheses.

We then used the sample-clustering module to test whether user-defined gene sets can resolve sample identity. Fourteen marker genes separated erythrocyte progenitors and hepatocytes into distinct clusters, whereas lung alveolar and lung pleural epithelial samples formed a shared cluster (Fig. 4B; Methods). Ten marker genes resolved five closely related pancreatic cell types (acinar, ductal, alpha, beta and delta) despite their shared tissue origin (Fig. 4C; Methods). Because these gene sets were selected for cell-type specificity, separation is expected by construction; this analysis therefore illustrates the workflow available to users rather than benchmarking marker performance in an unbiased manner.

Finally, genome-browser views of the *CD3E*, *PRSS1* and *ALB* loci confirmed cell-type-restricted hypomethylation over the called marker regions in grouped WGBS profiles from the corresponding T-cell, pancreatic acinar cell and hepatocyte populations (Fig. 4D; Methods), with per-group sample sizes indicated, allowing visual assessment of both marker specificity and within-group variability.

Together, these modules close the loop from marker retrieval to sample-level evaluation: users can move from a gene or locus of interest to its methylation behavior across cell types and to the discrimination of their own samples, without leaving the platform. This integrated functionality enables researchers to evaluate the biological relevance and discriminatory potential of DNA methylation markers, supporting their application in cell identity characterization and downstream methylome analyses.

### Cell-type identity recovery in an independent single-cell methylome atlas

To examine whether methylation-derived markers provide annotation signals that align more directly with single-cell methylome data than expression-derived marker genes, we mapped CellMethMarker custom markers to an independent human single-cell methylome atlas comprising 86,689 nuclei from 35 major cell types (Zhou et al. 2026) and scored each marker set by its median z-score in every atlas cell type (Fig. 5A; Methods). Based on cell-type identity, 24 of the 34 WGBS-derived and 17 of the 27 RRBS-derived custom marker cell types could be matched to a corresponding atlas cell type.

**Fig. 5:**
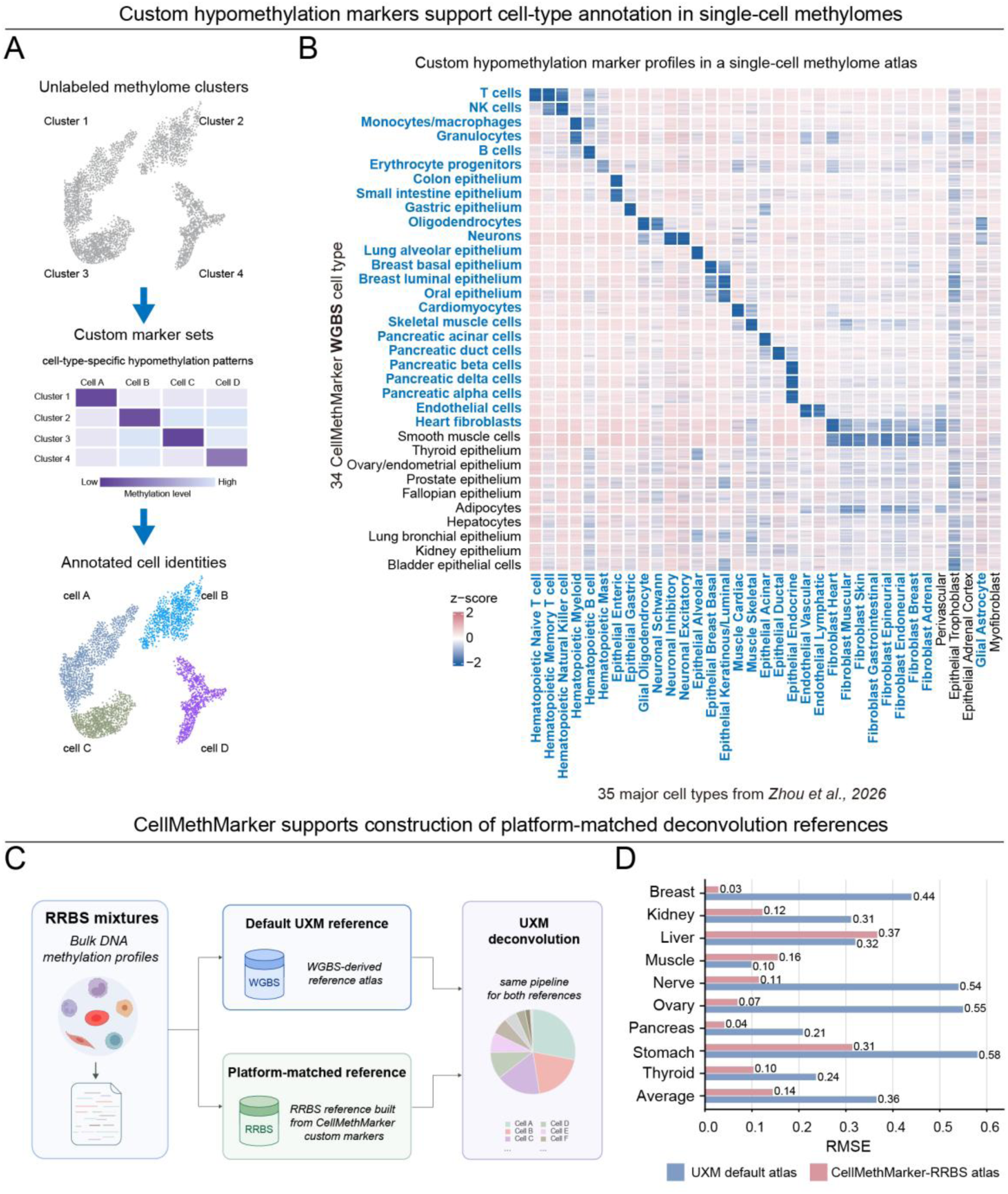
Case studies illustrating the utility of CellMethMarker custom markers in cell-type annotation and methylome deconvolution. **A** Schematic overview of the annotation strategy. Unlabeled methylome clusters are annotated by comparing their hypomethylation patterns with CellMethMarker custom marker sets, and cluster identities are assigned according to the best-matching cell type. **B** Row-wise z-scored methylation profiles of custom hypomethylated marker sets for 34 WGBS cell or tissue types across the 35 major cell types of the independent single-cell methylome atlas. Blue row and column labels indicate cell or tissue types matched between CellMethMarker and the atlas, whereas unmatched types are shown in black. Bold blue labels indicate matched types satisfying the predefined criterion for relative hypomethylation in the corresponding atlas cell type. Negative and positive *z* scores indicate methylation levels below and above the corresponding row mean, respectively, and their absolute values indicate the distance from the row mean in units of standard deviation. **C** Schematic overview of the benchmark. Identical RRBS mixtures were deconvolved using the same UXM pipeline with either the default WGBS-derived UXM reference or an RRBS-matched reference constructed from CellMethMarker custom markers. **D** Root mean square error (RMSE) between estimated and ground-truth proportions for nine benchmark cell and tissue types and the average across these types. Blue bars indicate results obtained with the default UXM atlas, and red bars indicate results obtained with the CellMethMarker-derived RRBS atlas. Lower RMSE indicates better deconvolution performance.

Under the predefined Top-1 and Top-3 recovery criteria (Methods), all 24 matched WGBS marker sets recovered their expected atlas cell types under both criteria (24/24, 100% for each; Fig. 5B and Supplementary Table S1), whereas 12 of the 17 matched RRBS marker sets achieved Top-1 recovery (12/17, 70.6%), and 15 achieved Top-3 recovery (15/17, 88.2%; Supplementary Fig. S2 and Supplementary Table S2). The recovered types span hematopoietic, epithelial, neural, muscle, pancreatic, endothelial and fibroblast lineages, indicating that the signal is not confined to the well-characterized blood compartment.

As a comparison matched in measurement context, we scored promoter methylation profiles of expression-derived marker genes from CellMarker 2.0 in the same atlas. A total of 57 CellMarker 2.0 cell-type marker sets could be mapped to 32 of the 35 distinct atlas cell types. Of these 57 marker sets, 18 achieved Top-1 recovery (18/57, 31.6%), collectively representing 14 distinct atlas cell types, and 30 achieved Top-3 recovery (30/57, 52.6%; Supplementary Fig. S3 and Supplementary Table S3). Thus, when the input data are themselves methylomes, region-level methylation markers provide a more directly matched annotation signal than promoter methylation of RNA-defined marker genes.

### A platform-matched reference improves cell-type deconvolution

The second case study asks a complementary question: whether a reference built from CellMethMarker markers improves a downstream quantitative task. Because RRBS measures only a small, CpG-island-biased fraction of the genome, many markers of a WGBS-derived reference are simply unmeasured in RRBS data, so platform mismatch — rather than the deconvolution model itself — may dominate the error. To test this, we deconvolved identical simulated RRBS mixtures with the same UXM pipeline, using either the default WGBS-derived UXM reference (Loyfer et al. 2023) or an RRBS-matched reference built from CellMethMarker custom markers with the top 25 markers per cell type (Fig. 5C; Methods). The RRBS-matched reference lowered the root mean square error (RMSE) for seven of nine benchmark cell and tissue types and reduced the mean RMSE from 0.365 to 0.144, a 60.5% reduction (Fig. 5D). Gains were largest where the default reference performed worst — ovary (0.55 to 0.07), nerve (0.54 to 0.11) and breast (0.44 to 0.03). Liver and muscle did not improve, indicating that performance remains bounded by cell-type representation, marker coverage and reference quality. Under the evaluated benchmark, these results indicate that deconvolution references should be matched to the assay — in chemistry and CpG coverage — and not only in biological content, a principle that CellMethMarker makes straightforward to implement.

## Discussion

A central challenge in methylome interpretation is determining whether a differential signal marks a stable cellular identity, reflects shifting sample composition, or simply annotates a biologically interesting genomic region. CellMethMarker addresses this challenge by treating cell- and tissue-type-specific DNA methylation markers as first-class, region-level objects rather than as auxiliary outputs of individual studies. By integrating uniformly derived WGBS and RRBS markers, manually curated literature records and harmonized sample metadata, the resource connects marker loci to the experimental context in which they were discovered and to the downstream settings in which they are most likely to be used. This complements disease-focused methylation databases, broad methylome atlases and RNA-centric cell-marker resources (Liu et al. 2016; Lebrón et al. 2017; Ding et al. 2020; Huang et al. 2021; Zong et al. 2022; Xing et al. 2022; Zhang et al. 2023; Hu et al. 2023; Jiang et al. 2023; Zhu et al. 2024), while filling a distinct niche: cross-study retrieval of human methylation markers whose primary annotation is cellular origin.

A distinctive feature of CellMethMarker is the emphasis on deeply curated, sorted or dissected discovery samples for *de novo* marker calling. This design increases the likelihood that the identified DMBs reflect cell- and tissue-intrinsic methylation programs rather than incidental variation in bulk-sample composition. The predominance of hypomethylated markers (94.9%) is also biologically consistent with previous human methylome atlases, in which loss of methylation often marks accessible, lineage-associated regulatory regions (Moss et al. 2018; Loyfer et al. 2023), and it carries a practical consequence: hypomethylation is the signal to look for when annotating an unknown methylome. Complementing the *de novo* layer, the literature-derived collection preserves study-specific provenance and captures marker types that are not uniformly represented in WGBS and RRBS block calls, including array-derived DMPs and DMRs. The joint availability of these layers allows users to distinguish markers supported by convergent evidence from those reported by a single platform or publication.

The two case studies illustrate how these markers can be used beyond static browsing. In the single-cell methylome analysis, CellMethMarker hypomethylated regions recovered the expected matched cell-type pattern more consistently than promoter methylation profiles derived from RNA-based marker genes under the same evaluation (24/24 WGBS-derived sets versus 18/57 CellMarker 2.0 marker sets). This does not imply that methylation markers should replace expression markers; rather, it shows that region-level methylation markers provide a more direct annotation signal when the input data are themselves methylomes. In the deconvolution benchmark, replacing the default WGBS-derived UXM reference with an RRBS-matched CellMethMarker reference reduced the mean RMSE from 0.365 to 0.144 under identical mixtures and model settings. The improvement supports a practical point for epigenetic liquid-biopsy and bulk-tissue analyses: reference construction should be matched not only to biological content but also to assay platform and CpG coverage.

These results should nevertheless be interpreted within clear boundaries. The annotation case study depends on the quality of cell-type matching between CellMethMarker and the independent atlas, and the CellMarker 2.0 comparison evaluates promoter methylation of marker genes rather than the full transcriptional program. The deconvolution analysis used simulated two-component mixtures across nine benchmark cell and tissue types and therefore cannot capture all complexities of clinical cell-free DNA, tumor heterogeneity or rare-cell detection; liver and muscle did not improve, indicating that marker coverage, reference completeness and biological similarity between components remain limiting factors.

Coverage of the resource is bounded by what is publicly available: cell types profiled by few or no public WGBS or RRBS datasets are correspondingly under-represented, and blood-derived samples account for nearly half of the metadata layer. Because *de novo* markers were called with a one-versus-rest design, their specificity is defined relative to the panel of cell types included and may weaken as the panel grows. Literature-derived records were not re-evaluated against raw data and should therefore be treated as curated evidence rather than as validated markers. More broadly, CellMethMarker is currently human-centered and hg19-centric for online search because most literature records were reported in hg19; although an hg38 liftOver version is provided, liftOver cannot fully recover regions that are unstable or unmappable between assemblies.

Future development will focus on four priorities. First, we will expand sorted-cell and dissected-tissue coverage, particularly for underrepresented tissues, developmental stages, immune subsets and disease-adjacent states. Second, we will incorporate additional platforms, including single-cell methylomes and array-based resources, where they can be harmonized without obscuring platform-specific resolution. Third, we will refine ontology mappings as CL, UBERON and MeSH evolve, and expose clearer provenance for terms that remain ambiguous. Fourth, we will strengthen programmatic access and benchmarking so that marker sets can be versioned, compared across releases and evaluated against community reference datasets. We are committed to maintaining CellMethMarker as a long-term community resource, with regular content updates and a submission channel for newly published marker sets. These updates should make CellMethMarker more useful for annotating single-cell methylomes, interpreting bulk and cell-free methylomes, and building platform-aware references for cell-origin analysis.

## Methods

### Database implementation

CellMethMarker was implemented using a decoupled web architecture. Django (v3.2.13; https://www.djangoproject.com/) provides the application and representational state transfer application programming interface (REST API) layer. MySQL (v8.0.42; https://www.mysql.com/) serves as the primary relational database for marker, sample, ontology and publication records, whereas Redis (v6.2.18; https://redis.io/) caches frequently accessed query results. The user interface was developed using HTML5, Bootstrap (v5.3.0; https://getbootstrap.com/), Tailwind CSS (https://tailwindcss.com/), and jQuery (v3.6.0; https://jquery.com/), with interactive charts rendered using amCharts 5 (https://www.amcharts.com/) and ECharts (Li et al. 2018). JBrowse 2 provides genomic visualization (Diesh et al. 2023). The frontend, backend, MySQL and Redis services are deployed in separate Docker containers (https://www.docker.com/) and orchestrated with Docker Compose (https://docs.docker.com/compose/). NGINX (https://nginx.org/) acts as the reverse proxy, and persistent database and genome-browser files are stored in mounted volumes.

### Literature marker collection and curation

All cell- or tissue-specific DNA methylation markers from the literature were manually extracted from publications through May 2025 indexed in the PubMed database. We searched PubMed for relevant studies using a list of keywords appearing in the title, abstract, or full text, such as “cell-type specific”, “tissue-specific”, “DNA methylation” and “differential methylation”. A total of 733 publications were initially collected. We then manually screened each article, excluding studies that (i) were not based on human samples or (ii) did not identify any markers through differential methylation analysis between cell or tissue types. After this screening process, 78 publications were retained.

Marker records were extracted from the main text, supplementary materials and associated repositories, and standardized to a uniform format. Identical loci reported across different studies were retained as independent evidence records to preserve study-specific provenance. To harmonize genome assemblies, genomic coordinates were converted either to hg19 (from hg18 or hg38) or to hg38 (from hg18 or hg19) using the liftOver function implemented in the rtracklayer R package (Lawrence et al. 2009). Relative to all 2,203,507 curated records, 658 coordinates (0.030%) failed to map to hg19 and 541 (0.025%) failed to map to hg38; coordinates that could not be successfully mapped to the target genome assembly were excluded from the corresponding release. Functional annotations were performed using ChIPseeker (Yu et al. 2015) and annotatr (Cavalcante and Sartor 2017). In total, we curated 2,203,507 methylation markers, which are documented as “Literature Markers” in CellMethMarker.

### Sample metadata collection

A total of 2,810 samples were included, of which 2,762 were retrieved from the NCBI Gene Expression Omnibus (GEO; https://www.ncbi.nlm.nih.gov/geo/) (Barrett et al. 2013). The remaining 48 samples were generated as part of a separate ongoing study within our research group and were made available for secondary analysis in the present study; these data have been deposited in the Open Archive for Miscellaneous Data (OMIX; https://ngdc.cncb.ac.cn/omix/) (Zhang et al. 2025; CNCB-NGDC Members and Partners 2026) database under accession OMIX012321. No new human samples were collected specifically for the present study. The curated metadata include cell or tissue type, ontology identifier, sex, age, cell-sorting strategy, medical condition and DNA methylation assay.

### Ontology harmonization

Cell- and tissue-type annotations were harmonized using the Cell Ontology (CL; release 2026.6.8) (Diehl et al. 2016), the Uber-anatomy ontology (UBERON; release 2026.4.1) (Mungall et al. 2012) and Medical Subject Headings (MeSH; version 2025 as indexed in Ontology Lookup Service, OLS4, accessed 23 June 2026). CL and UBERON ontology files were downloaded in Open Biomedical Ontologies (OBO) format on 23 June 2026 and parsed locally. Source labels were normalized by conversion to lowercase, standardization of punctuation and non-alphanumeric characters, and normalization of common singular and plural forms. The normalized labels were first matched against ontology preferred labels and exact synonyms. Cell-type terms were preferentially mapped to CL, whereas tissue and anatomical terms were preferentially mapped to UBERON.

Ambiguous matches, labels associated with multiple candidate terms and labels that could not be mapped automatically were manually reviewed using OLS4. When no suitable CL or UBERON term was available, the most specific semantically appropriate MeSH descriptor consistent with the original label, biological context and source metadata was selected. For each confidently mapped entry, the source vocabulary, accession identifier and preferred label were recorded. Original standardized labels were retained for terms that could not be mapped with sufficient confidence, and the corresponding ontology fields were left blank.

### Processing of methylation profiles and genome browser integration

We curated high-quality DNA methylation profiles for 2,171 samples, comprising 1,137 RRBS, 988 WGBS, 10 eRRBS and 36 mRRBS samples. Raw sequencing reads in FASTQ format for 667 RRBS and 264 WGBS samples were downloaded from GEO, the Sequence Read Archive (SRA; https://www.ncbi.nlm.nih.gov/sra/) (Kodama et al. 2012), the European Nucleotide Archive (ENA; https://www.ebi.ac.uk/ena/) (Yuan et al. 2026), and the European Genome-phenome Archive (EGA; https://ega-archive.org/) (Lappalainen et al. 2015) and processed using our customized analysis pipeline described in the “Custom marker data processing” section below.

The remaining processed methylation profiles were retrieved from GEO. Because these profiles were generated using different study-specific processing pipelines, we inspected the file structure and methylation reporting format of each dataset before subsequent harmonization. To standardize the datasets, we employed custom format conversion scripts, resulting in a consistent five-column format: chr start end meth total. For datasets aligned to the hg38/GRCh38 reference genome, we utilized the ucsc-liftover (v469) tool to convert coordinates to hg19/GRCh37, discarding loci that could not be mapped.

All 2,171 methylation profiles were converted to BigWig format and integrated into the genome browser. For cell- and tissue-type-level visualization, the profiles were summarized in 200-bp windows using deepTools (Ramírez et al. 2016) and grouped by methylation assay and standardized cell or tissue type, enabling both individual-sample and grouped visualization across RRBS, WGBS, eRRBS and mRRBS datasets. The resulting aggregated tracks, together with the individual-sample tracks, were used to construct the genome browser.

### Custom marker data processing

#### RRBS raw sequencing reads processing

Raw FASTQ data for all 667 RRBS samples were retrieved from the ENA website, comprising 419 single-end and 248 paired-end samples. Data processing was performed to capture methylation information, beginning with quality control using FastQC (v0.12.1). Adapter sequences were trimmed, and the data were cleaned using Trim Galore (v0.6.10). Alignment to the hg19 reference genome was conducted with BSMAP (v2.90) (Xi and Li 2009), followed by duplicate read marking using the markdup command in Sambamba (v1.0; Tarasov et al. 2015). Methylation information was then extracted using MethylDackel (v0.6.1).

To ensure sufficient data quality and CpG coverage for downstream analyses, we first assessed CpG sites covered at ≥10× depth across all RRBS samples. Samples with fewer than 1,000,000 CpG sites meeting this criterion were excluded, resulting in removal of 66 samples. Additional exclusions were applied based on biological relevance and experimental consistency. Specifically, we excluded samples derived from cell lines (n = 3), non-neuronal cells (n = 6), and induced pluripotent stem cells (iPSCs; n = 2), as these samples were either reprogrammed or not directly purified from primary tissues. Hematopoietic stem cell (HSC) samples with insufficient marker availability ( ≤ 3 analyzable markers; n = 6) were also excluded. Moreover, T cell samples treated with 5-aza-2′-deoxycytidine (AZA-CdR; n = 3) and CD3+ T cell samples from post-treatment diffuse large B-cell lymphoma (DLBCL) patients that originated from mixed tissue and peripheral blood sources (n = 6) were removed.

After quality control and biological filtering, 575 RRBS samples were retained. These samples were combined with 521 tissue RRBS samples from GSE233417 (Li et al. 2023), for which raw sequencing reads were unavailable but processed methylation data in .pat format were accessible for downstream analysis, and 48 in-house normal tissue RRBS samples, yielding a total of 1,144 samples. Samples were grouped according to their annotated cell or tissue types for subsequent analyses. To reduce sampling bias and ensure group comparability, white blood cell (WBC) samples were removed from GSE233417, and the remaining samples were manually downsampled to retain 3–36 samples per group, without applying a fixed computational randomization procedure. Differential methylation analysis was then performed iteratively using alternative sample subsets and cell/tissue group compositions. Group inclusion and sample selection were refined through empirical assessment of marker yield, cell/tissue-type coverage, data quality, and the biological consistency of the resulting marker sets. Groups with insufficient marker representation, limited sample availability, or questionable biological assignments were excluded during this process. Ultimately, 367 RRBS samples were retained for marker discovery. The identifiers and corresponding annotations of the final discovery samples are provided in Supplementary Table S4.

#### WGBS raw sequencing reads processing

For WGBS samples with downloadable raw sequencing reads, we retrieved raw FASTQ data from the EGA and ENA repositories. Data processing was conducted using a custom pipeline, based on the CpG Me approach (Krueger and Andrews 2011; Martin 2011; Ewels et al. 2016; Laufer et al. 2022), which incorporates a WGBS alignment pipeline utilizing the hg19/GRCh37 reference genome. We then implemented a multi-step filtering strategy based on sequencing quality, cellular group composition, and biological relevance. First, to ensure sufficient and comparable coverage across samples, we excluded datasets with excessive missing data (>80% missing CpG measurements), which were predominantly generated on lower-throughput sequencing platforms. Samples lacking sufficient coverage at shared CpG markers were deemed unsuitable for combined differential methylation analysis and were removed. Second, cell types represented by a single sample or by overly heterogeneous processing pipelines were excluded. Third, samples derived from cell lines, reprogrammed cells, or specialized experimental manipulations were excluded. In addition, samples lacking identifiable differential methylation markers or representing closely related developmental cell states that could not be reliably distinguished were removed. After applying these filtering steps, a high-quality and biologically consistent subset of 205 WGBS samples was retained for custom marker identification. The identifiers and corresponding annotations of the final discovery samples are provided in Supplementary Table S4.

#### De novo discovery of custom markers

Custom markers were identified separately for WGBS and RRBS data. Raw sequencing data were processed using standardized, platform-specific pipelines, while externally processed methylation profiles were harmonized into a common format before marker discovery. Samples from different datasets and sources were combined and grouped by their annotated cell or tissue types. Differential methylation analysis was then performed using a one-versus-rest strategy, comparing each cell or tissue type with all remaining types to identify cell- or tissue-type-specific differentially methylated blocks (DMBs). The genome was first segmented into homogeneously methylated blocks using the wgbstools (Loyfer et al. 2026) “segment” subcommand, after which the “beta_to_table” subcommand was used to aggregate beta values within each block. Cell-type-specific DMBs were then identified using the wgbstools “find_markers” subcommand with pval = 0.05. For RRBS data, the parameters were set to min_cpg = 2, max_bp = 1500, delta_quants = 0.1, and min_cov = 3, whereas for WGBS data, min_cpg = 4, max_bp = 1500, and delta_quants = 0.2 were used. This workflow identified a total of 347,514 cell-type-specific DMBs, which were incorporated into CellMethMarker.

### Classification into broad biological categories

For visualization and comparative analysis, standardized cell and tissue types were grouped into nine broad biological categories: hematopoietic and immune; nervous and sensory; cardiorespiratory; digestive and endocrine; urogenital and reproductive; integumentary and mammary; musculoskeletal, stromal and adipose; pluripotent and developmental; and mixed and unspecified targets. Classification was guided by cellular lineage, anatomical origin, and physiological function, drawing on the organizational principles of the Cell Ontology (Diehl et al. 2016), UBERON (Mungall et al. 2012), and Human Cell Atlas (Regev et al. 2017). Cell types were assigned according to their predominant lineage or anatomical context, with pluripotent and developmental populations categorized separately. Composite or insufficiently specified targets were assigned to the mixed and unspecified category. These broad categories were defined for descriptive analysis rather than as formal ontology classes. Original annotations were preserved, and complete assignments are provided in Supplementary Table S5.

### Web-based marker analysis and visualization

Methylation heatmap visualization, sample clustering, and genome-browser inspection were performed using the integrated web modules of CellMethMarker. For the methylation heatmap, a panel of 60 canonical cell-identity genes was submitted to the “Methylation Heatmap” module (Supplementary Table S6). WGBS was selected as the methylation assay, and “Column clustering” was enabled under “Clustering Options”. Mean methylation levels of gene-associated custom marker regions were visualized across cell and tissue types.

For sample clustering, two gene panels comprising 14 marker genes representing erythrocyte progenitors, hepatocytes, lung alveolar epithelium, and lung pleural epithelium (Supplementary Table S7) and 10 marker genes representing five pancreatic cell types (Supplementary Table S8) were analyzed separately using the “Sample Clustering” module. For both analyses, WGBS was selected as the methylation assay, “Feature level” was set to “gene-level average”, and “Hierarchical clustering” was selected as the clustering algorithm. The number of clusters was set to four for the diverse cell-type analysis and six for the pancreatic cell-type analysis. Gene-level methylation features were calculated by averaging methylation levels across the marker regions associated with each gene. Features with valid methylation measurements in fewer than 5% of samples (or fewer than two samples) were excluded, and samples with valid measurements for fewer than 30% of the retained features were removed. Missing values were imputed using feature-wise means, followed by Z-score standardization. Agglomerative hierarchical clustering was performed using Euclidean distances between cluster centroids. Principal component analysis (PCA) was independently applied to the standardized feature matrix, and the first two principal components were used as the coordinates for two-dimensional visualization of the clustering results.

Finally, the “Genome Browser” module was used to inspect the *CD3E*, *PRSS1*, and *ALB* loci, displaying gene annotations, custom marker regions, and cell/tissue-grouped WGBS methylation profiles in the hg19 reference genome. The corresponding tracks were selected from “Methylation Profiles” > “WGBS” > “Cell-type Grouped Sample Profiles” in the “Available tracks” panel, with “xyplot” selected as the renderer type. For *CD3E*, the selected tracks included CD4+ regulatory T cells (n = 10), CD4+ conventional T cells (n = 7), breast basal epithelium (n = 4), and hepatocytes (n = 6). For *PRSS1*, the selected tracks included pancreatic acinar cells (n = 4), pancreatic alpha cells (n = 3), pancreatic beta cells (n = 3), pancreatic delta cells (n = 3), pancreatic duct cells (n = 4), and hepatocytes (n = 6). For *ALB*, the selected tracks included hepatocytes (n = 6), breast basal epithelium (n = 4), CD4+ T cells (n = 23), and colon epithelium (n = 5).

### Cell-type annotation case study

We first obtained single-nucleus methyl-3C sequencing (snm3C-seq) methylation profiles for 35 major cell types (Zhou et al. 2026). Human cell markers were downloaded from the CellMarker 2.0 database (Hu et al. 2023) and filtered according to the following criteria: species = Human, cancer type = Normal, and cell type = Normal cell. After harmonizing cell-type annotations between CellMarker 2.0 and the snm3C-seq atlas, 57 CellMarker 2.0 cell types were matched to 32 of the 35 atlas cell types. For each CellMarker 2.0 cell type, up to five canonical marker genes were manually selected from the retrieved records based on published literature supporting their use as markers of the corresponding cell type. Marker selection was performed independently of methylation patterns in the evaluation atlas. Promoter regions were defined as the intervals spanning 2,000 bp upstream to 500 bp downstream of the transcription start site.

We next selected cell-type-specific hypomethylated markers from the CellMethMarker custom marker collection. Among the RRBS-derived custom markers, 17 of the 27 represented cell types could be matched to 29 of the 35 cell types in the snm3C-seq atlas. Similarly, among the WGBS-derived custom markers, 24 of the 34 represented cell types could be matched to 31 atlas cell types. For each matched CellMethMarker cell type, up to 250 hypomethylated markers with the greatest differential methylation magnitude were retained. Finally, bigWigAverageOverBed was used to calculate methylation levels across the 35 major atlas cell types within the selected CellMethMarker hypomethylated marker regions and within the promoter regions of the CellMarker 2.0 marker genes.

For each marker region, methylation values were standardized across the 35 major cell types in the Zhou et al. single-cell methylome atlas using a row-wise z-score transformation. Specifically, the z-score for each marker in a given atlas cell type was calculated relative to the mean and standard deviation of that marker across the available atlas cell types. Marker regions with methylation measurements available in fewer than five atlas cell types were excluded before standardization, and undefined z-scores were set to zero. The standardized marker profiles were then grouped according to their corresponding CellMethMarker or CellMarker 2.0 cell types. For each marker set, the median z-score across all constituent markers was calculated separately for each of the 35 atlas cell types, with lower values indicating stronger relative hypomethylation. The atlas cell types were subsequently ranked according to these median z-scores. A marker set was considered successfully recovered under the Top-1 criterion when its predefined corresponding atlas cell type exhibited the lowest median z-score among all 35 atlas cell types. Top-3 recovery was additionally recorded when the corresponding atlas cell type ranked among the three lowest median z-scores.

### Cell-type deconvolution case study

We selected nine RRBS samples representing breast, kidney, liver, muscle, nerve, ovary, pancreas, stomach and thyroid tissues or cell types. None of these nine target samples was used for custom marker discovery or construction of the CellMethMarker-RRBS reference atlas. Each target sample was mixed *in silico* with a leukocyte RRBS sample using wgbstools “mix_pat” subcommand. Simulated mixtures were generated through random fragment sampling, with random seeds specified to ensure reproducibility. The target-sample fraction was varied from 0 to 1 in increments of 0.1, with the leukocyte fraction adjusted accordingly so that the two components summed to 1. Ten replicate mixtures were generated independently at each mixing proportion using different random seeds, yielding 110 mixtures per target and 990 simulated mixtures in total. Because the default UXM reference atlas contains 25 markers per cell type, we selected up to 25 cell-type-specific hypomethylated markers with the largest differential methylation values for each cell type from the CellMethMarker custom marker collection. These markers were used with the UXM build function to construct an RRBS-matched reference atlas, hereafter referred to as the CellMethMarker-RRBS atlas. The simulated mixtures were then deconvolved with UXM using either the default reference atlas or the CellMethMarker-RRBS atlas under otherwise identical settings. Deconvolution accuracy was assessed for each cell or tissue type by calculating the root mean square error (RMSE) between the estimated and ground-truth proportions.

### Statistics and Reproducibility

CellMethMarker is a curated resource, and the descriptive statistics reported in Fig. 2 are counts and percentages computed over the complete curated collections rather than over samples drawn from them; no hypothesis tests were applied to the database content. Differential methylation strength is summarized as the mean and median Δβ across all markers of a given assay, with violin plots showing the full distribution and boxplots showing the median, interquartile range and 1.5 times the interquartile range. Marker discovery used a one-versus-rest design after sample quality control, as described above. In the cell-type annotation case study, methylation values for each individual marker region were standardized across atlas cell types using a row-wise z-score transformation. For each marker set, the median of the constituent marker z-scores was then calculated for each of the 35 atlas cell types and used to rank them by relative hypomethylation. The Top-1 and Top-3 recovery criteria described above were applied consistently to WGBS-derived marker sets, RRBS-derived marker sets and CellMarker 2.0 promoter marker sets. The final sample sets used for *de novo* marker discovery were determined through iterative, empirically guided selection, and their complete identifiers are provided in Supplementary Table S4. In the deconvolution benchmark, ten replicate *in silico* mixtures were generated at each of 11 mixing proportions for each of nine target cell or tissue types (110 mixtures per target, 990 in total), using specified random seeds to enable reproducible simulation. Both references were evaluated on exactly the same mixtures with identical UXM settings, and accuracy was summarized as the RMSE between estimated and ground-truth proportions across the 110 mixtures of each target. No additional data were excluded after applying the predefined quality-control, coverage and marker-availability filters described above, and randomization and blinding were not applicable. Software versions, reference assemblies and key parameters are provided in the relevant Methods subsections.

## Data access

CellMethMarker is publicly accessible at http://gu-zhanglab.hfcas.ac.cn/CellMethMarker and requires no registration. Custom markers, literature-derived markers in hg19 and hg38, harmonized sample metadata and sample-level BigWig methylation profiles are available in full through the database search and download interfaces. The 48 in-house samples are deposited in the OMIX database under accession OMIX012321 (https://ngdc.cncb.ac.cn/omix/preview/hhdkxDTLUU). Publicly available datasets reanalyzed in this study were obtained from GEO, SRA, ENA and EGA under the accessions listed in Supplementary Table S4. The single-cell methylome atlas used in the annotation case study is available from the original publication (Zhou et al. 2026).

## Acknowledgements

We thank the investigators and consortia who generated and shared the public methylation datasets used in this study. We gratefully acknowledge the members of the Epigenomics and Intelligent Analysis team for their assistance with the collection and curation of publicly available datasets used in this study. Some graphical elements in Figs. 1 and 5C were obtained from BioRender.com and assembled by the authors. This work was supported by the Key Science & Technology Project of Anhui Province (Grant No. 202523n10050006 to F.Z.), the Anhui Provincial Natural Science Foundation (Grant No. 2408085J017 to F.Z.), the Hundred Talents Program to F.Z., the Noncommunicable Chronic Diseases-National Science and Technology Major Project (Grant No. 2024ZD0520100 to H.G.) and the CASHIPS Seed Grant to H.G. F.Z., J.S., H.W., and H.G. supervised the project. F.Z. and X.S. conceived and designed the study. X.S., Y.P., and Y.Y. developed the database and website, designed and implemented the web interface, optimized the page layout and styling, and performed website testing and quality control. X.S., L.F., F.L. and Y.X. curated the data; X.S. performed the data analyses. X.S. wrote the original draft of the manuscript. J.S., F.Z., H.G., and H.W. reviewed and edited the manuscript and provided critical feedback throughout the project. All authors reviewed and approved the final manuscript.

## Software availability

The custom pipelines for methylation profile processing, *de novo* marker discovery, and the case study are available on Zenodo (https://doi.org/10.5281/zenodo.22012363). Third-party software, versions and non-default parameters used to process methylation profiles, construct marker sets and implement the website are reported in the Methods.

