## Supplemental figures for "CellMethMarker provides an integrated database of human cell- and tissue-type-specific DNA methylation markers"

**Contents:**

Supplementary Figures

Supplementary Fig. S1

Supplementary Fig. S2

Supplementary Fig. S3

### Supplementary Figures

#### Supplementary Fig. S1

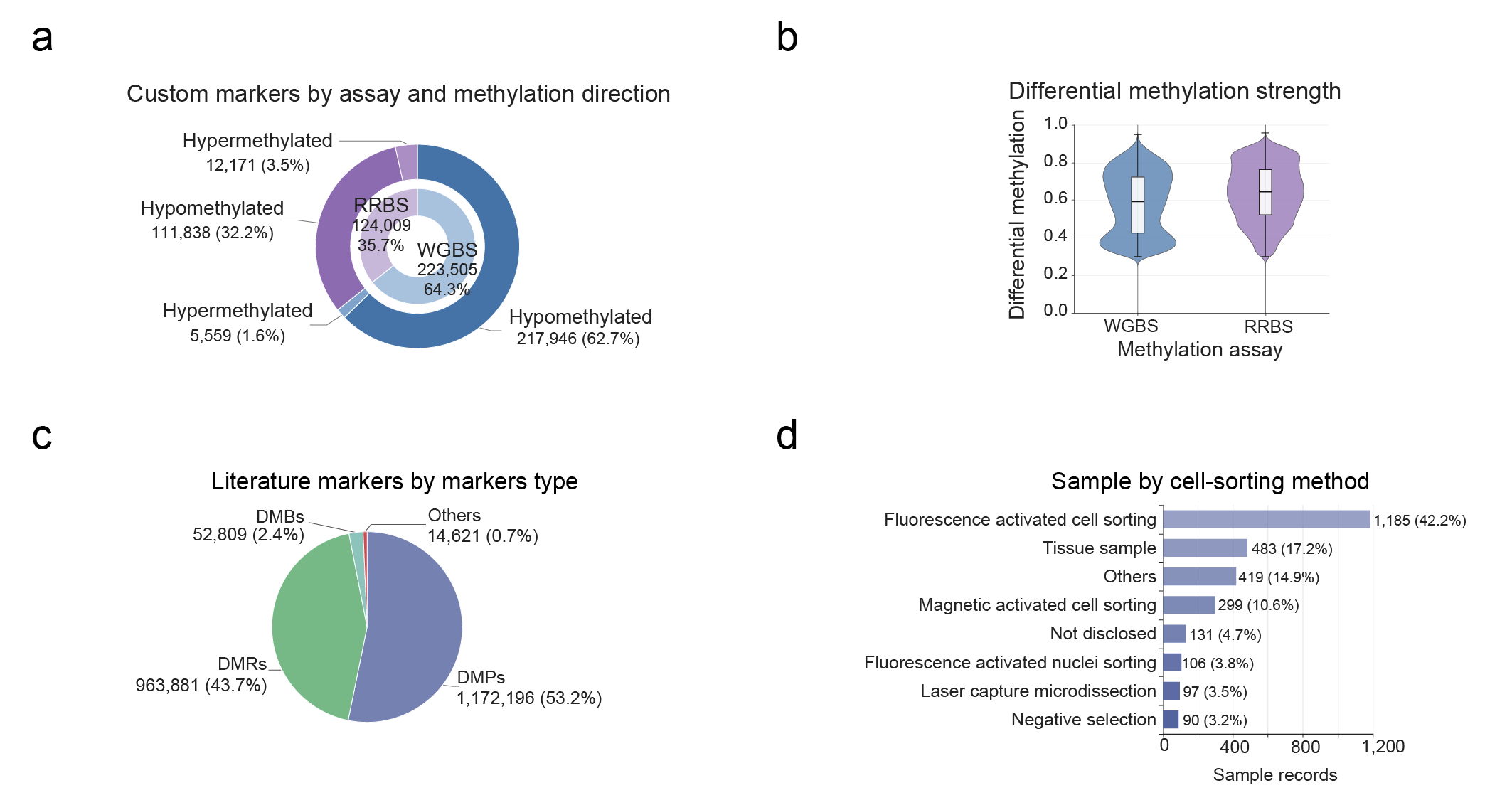

**Supplementary Fig. S1 | Coverage of the resource. a** Custom markers by methylation assay (inner ring) and methylation direction (outer ring). **b** Differential methylation strength (Δβ) of WGBS- and RRBS-derived markers. Violins show the full distribution; boxes show the median and interquartile range, and whiskers extend to 1.5 times the interquartile range. **c** Literature-derived marker records by marker type, marker types outside the categories shown are pooled as “Others”. **d** Curated samples by cell-sorting method. WGBS, whole-genome bisulfite sequencing; RRBS, reduced-representation bisulfite sequencing; DMP, differentially methylated position; DMR, differentially methylated region; DMB, differentially methylated block.

#### Supplementary Fig. S2

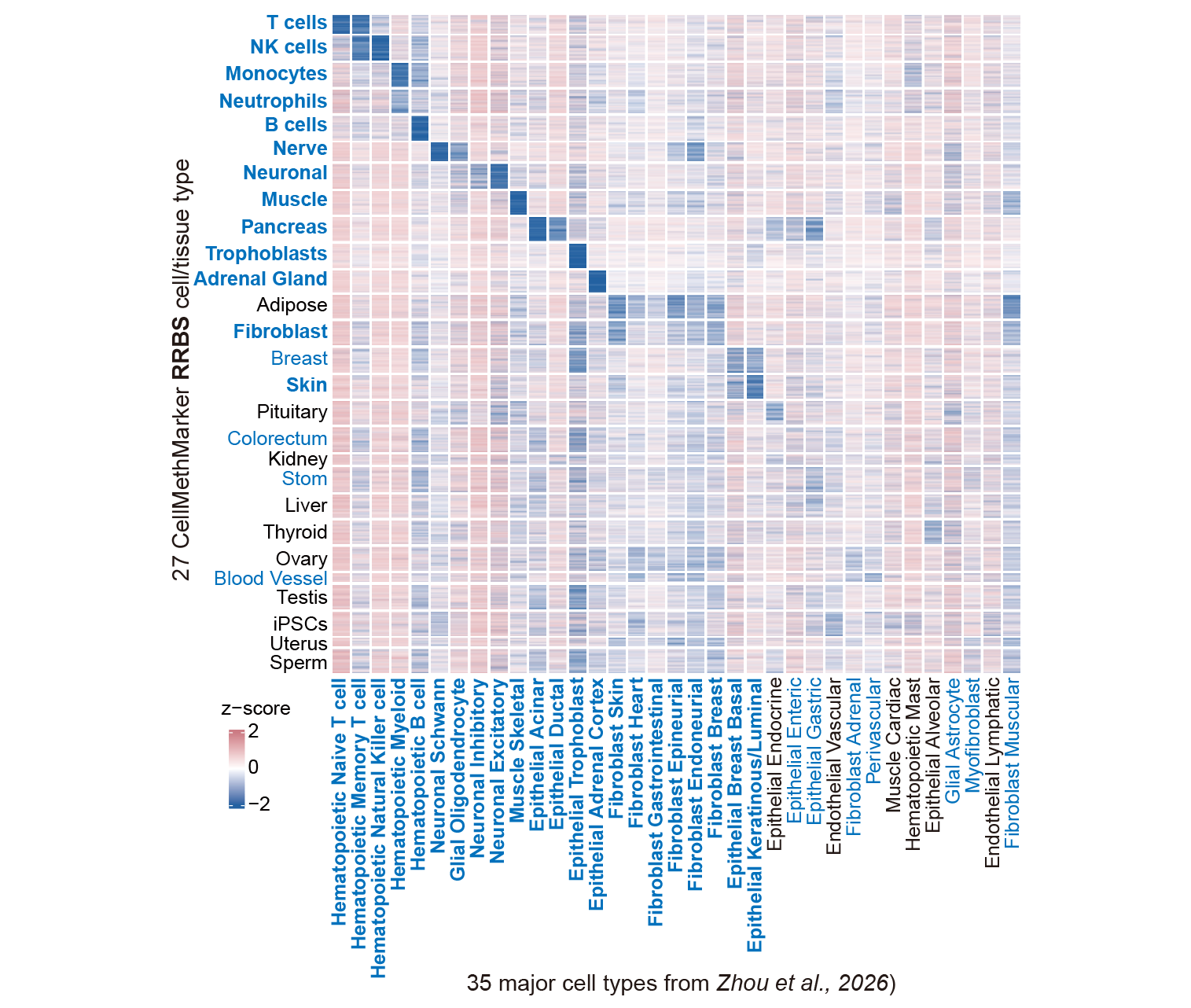

**Supplementary Fig. S1 | Annotation of an independent single-cell methylome atlas with CellMethMarker custom RRBS markers.** Row-wise z-scored methylation profiles of custom hypomethylated marker sets for 27 RRBS cell or tissue types (rows) across the 35 major cell types of the independent single-cell methylome atlas (columns). Blue row and column labels indicate cell or tissue types matched between CellMethMarker and the atlas, whereas unmatched types are shown in black. Bold blue labels indicate matched types satisfying the predefined criterion for relative hypomethylation in the corresponding atlas cell type. For each marker region, methylation values were standardized within each row across the 35 atlas cell types and are displayed as z scores. Negative and positive z scores indicate methylation levels below and above the corresponding row mean, respectively, and their absolute values indicate the distance from the row mean in units of standard deviation.

#### Supplementary Fig. S3

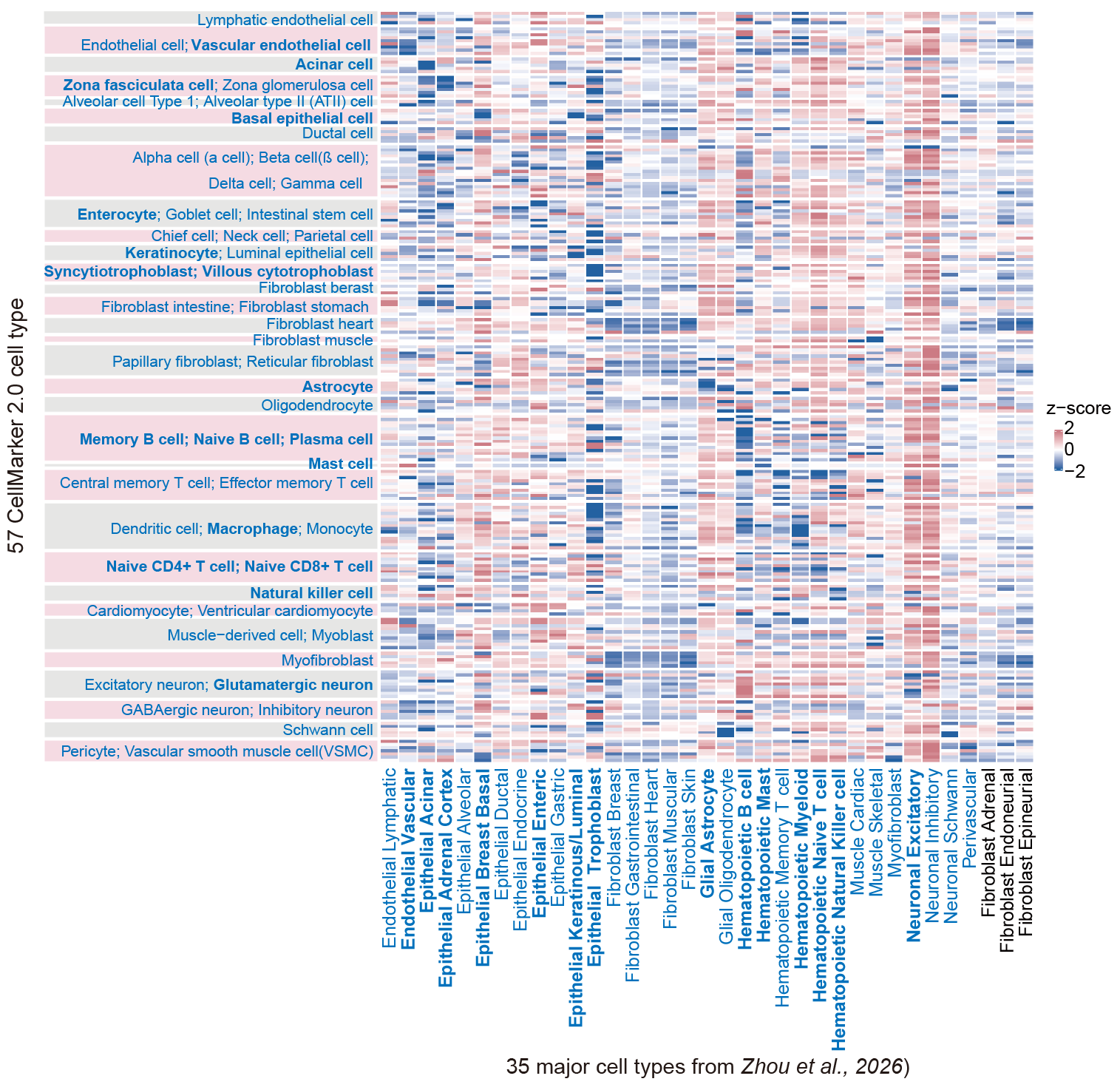

**Supplementary Fig. S2 | Promoter methylation patterns of CellMarker 2.0 marker genes across an independent single-cell methylome atlas.** The heatmap shows promoter methylation profiles of expression-derived marker genes from CellMarker 2.0 across 35 major cell types in the single-cell methylome atlas of Zhou *et al*. (2026). Rows represent 57 CellMarker 2.0 cell-type categories, with rows sharing the same background shading grouped according to their corresponding major cell type. Columns represent the 35 major cell types in the atlas. Blue row and column labels indicate cell or tissue types matched between CellMarker 2.0 and the atlas, whereas unmatched types are shown in black. Bold blue labels indicate matched types satisfying the predefined criterion for relative hypomethylation in the corresponding atlas cell type. Methylation values were standardized within each row across all atlas cell types and are displayed as z-scores. Negative and positive z-scores indicate methylation levels below and above the corresponding row mean, respectively, with their absolute values reflecting the magnitude of deviation in units of standard deviation.
